# Prevalence and correlates of anemia among children aged 6-23 months in Wolaita Zone, Southern Ethiopia

**DOI:** 10.1101/441196

**Authors:** Mihiretu Alemayehu, Mengistu Meskele, Bereket Alemayehu, Bereket Yakob

## Abstract

**Background:** Anemia, the world’s most common micro-nutrient deficiency disorder, can affect a person at any time and at all stages of life, although children aged 6 -23 months are particularly at higher risk. If left untreated, it adversely affects the health, cognitive development, school achievement, and work performance. However, littlewas investigated among young children in Sub-Saharan countries including Ethiopia. This research aimed to investigate its magnitude and correlates to address the gap and guide design of evidence based intervention.

**Methods:** A community-based cross-sectional study was conducted from May -June 2016 in rural districts of Wolaita Zone. Multi-stage sampling technique was applied to select 990 mother-child pairs. Socio-demography, health and nutritional characteristics were collected by administering interview type questionnaire to mothers/care-givers. Blood samples were taken to diagnose anemia by using HemoCue device, and status was determined using cut-offs used for children aged 6-59 months. Hemoglobin concentration below 11.0 g/dl was considered anemic. Data were analyzed with Statav 14. Bivariate and multivariable logistic regressions were applied to identify candidate and predictor variables respectively. Statistical significance was determined at p-value < 0.05 at 95% confidence interval.

**Results:** The mean hemoglobin level of children was 10.44±1.3g/dl, and 65.7% of them were anemic. Among anemic children, 0.4% were severely anemic (<7.0g/dl), while 28.1% and 37.2% were mildly (10.0-10.9g/dl) and moderately (7.0-9.9g/dl) anemic, respectively. In the multivariable analysis, having maternal age of 35 years and above (AOR=1.96), being government employee (AOR=0.29),being merchant (AOR= 0.43) and ‘other’ occupation (AOR=3.17) were correlated with anemia in children in rural Wolaita. Similarly, receiving antihelminthic drugs (AOR= 0.39), being female child (AOR= 1.76), consuming poor dietary diversity (AOR=1.40), and having moderate household food insecurity (AOR=1.72) were associated with anemia in rural Wolaita.

**Conclusion:** A large majority of children in the rural Wolaita were anemic and the need for proven public health interventions such as food diversification, provision of anti-helminthic drugs and ensuring household food security is crucial. In addition, educating women on nutrition and diet diversification, as well as helping them with alternative sources of income might be interventions in the study area.

## Background

Anemia is the world’s most common micro-nutrient deficiency disorder that affects more than 2 billion people globally [1]. It can affect a person at any time and at all stages of life. However, in most parts of the world, children aged 6-23months are at particularly higher risk. It primarily affects infants and young children because of their higher iron requirements related to growth, and women of childbearing age due to menstrual loss and pregnancy [2].

If left untreated, anemia can adversely affect the health, cognitive development, school achievement, and work performance of individuals. Low oxygenation of brain tissues, a consequence of anemia, may lead to impaired cognitive function, growth and psychomotor development in children. Infants, under 5-year-old children and pregnant women have greater susceptibility to anemia because of their increased iron requirements for rapid body growth and expansion of red blood cells [3]. During childhood period, it is strongly associated with poor health and physical development [4, 5], mild and moderate mental retardation [6], and poor motor development and its control [7]. Iron deficiency anemia leads to reduced academic achievement and work capacity which reduces the earning potential of individuals and hence damages national economic growth at large [5]. It also increases the risk of mortality and morbidity from infectious diseases [4, 8].

Childhood anemia is mostly caused by dietary iron deficiency, infectious and genetic diseases, and other nutrient deficiencies [9]. While more than half of the anemia burden in children is attributed to iron deficiency, only very small fraction is due to genetic causes [10]. In early childhood, bad feeding habits, especially during the weaning period as breast milk is replaced by foods that are poor in iron, vitamin B12 and folic acid [11].

In Ethiopia, evidence showed varying magnitudes of anemia ranging from 41% in Amhara region to 83% in Somali region [12]. Many studies showed that factors contributing to and affecting anemia in children varied from one geographical area to another [4, 6, 7, 10]. However, most of those studies were based on small sample size, conducted on urban dwellers, were facility-based, and lacked representativeness. Besides, no published research exists which uncovered the problem in the context of Woita Zone. Hence, the present study was conducted to fill the gap in a rural setting in Wolaita Zone with an aim of determining the magnitude of anemia and associated factors among children aged 6-23 months.

## Methodology

### Study design and Study area

A community based cross-sectional study was conducted among children aged 6-23 months residing in rural districts of Wolaita Zone, Southern Ethiopia, from May to June 2016. Wolaita Zone is one of the 13 administrative zones of southern region (SNNPR) which has 12 rural and 3 urban districts. The Zone was inhabited by over 1.8 million people in 2016 [13]. Wolaita Sodo, the capital of the Zone, is located at 6° 49’ N latitude and 39° 47’ E longitude, at an altitude of about 1900 meters above sea level. It is located at 330 km south-west of Addis Ababa, Ethiopia. The zone is characterized by its dense population. The majority (88.3%) of its population reside in rural districts whose major livelihood is agriculture. The major food crops cultivated in the zone are maize, sweet potato, *enset* (false banana), *teff* (*Eragrostistef*), haricot bean, taro, sorghum, Irish potato, yam and cassava [14].

### Sampling

Multi-stage sampling technique was applied to select mother-child paired study population. Children aged 6-23 months were the source population for the study. Initially, four districts were randomly selected from the 12 rural districts. Damot Gale, Boloso Bombe, Humbo and Sodo Zuria districts were selected as study districts. Then, three kebeles (the lowest administrative unit of Ethiopia consisting of nearly 5000 population) were randomly selected from each of the selected districts, making the total number of kebeles included in the study 12. Finally, the study participants were selected through systematic sampling technique from each of the selected kebeles by probability proportional to size i.e., allocating the sample size with regard to the respective kebeles’ population.

The total 993 sample size needed for this study was determined by the formula to estimate a single population proportion based on the following assumptions: 71.2% prevalence of anemia in Sub-Saharan Africa [15], 95% confidence interval, 5% margin of error, and 10% non-response rate and design effect of 3.

### Data collection tools and procedures

Data were collected through interviewer administered questionnaire prepared in English and translated to local language (Wolaita Dona). The questionnaire was developed by reviewing guidelines and related literatures [12, 15, 16, 17]. It consisted of demographic characteristics variables, household wealth indicators and anemia risk factors such as health service utilization, recent illnesses, and dietary practice of both mother and child.

#### Anemia diagnosis

Hemoglobin concentration was used to determine anemia status of the study participants by taking finger-prick blood sample. Hemoglobin level was analyzed onsite by using HemoCue device (HemoCueHb 301), and values were adjusted for altitude using the UNICEF/WHO guideline. Anemia status was determined using cut-offs used for children aged 6-59 months. Hemoglobin concentration below 11.0 g/dl was considered anemic, whereas, hemoglobin concentration of 11.0 g/dl and above was considered as normal. Severity of anemia was categorized based on the UNICEF, UNU, WHO guideline as follows: children were categorized as mildly, moderately and severely anemic if their blood hemoglobin concentration is between 10.0-10.9g/dl, 7.0-9.9g/dl and < 7.0 g/dl, respectively. Maternal anemia (hemoglobin concentration below 12 g/dl) was diagnosed with the same procedure and device used for child anemia, followed by adjustment for pregnancy and altitude [16].

#### Dietary assessment

A 24-hour dietary recall method was used to assess dietary practice. Dietary Diversity Score of children was calculated by asking mothers/caregivers about the food items their children consumed in the past 24 hours preceding the survey. All food items consumed by the children in the last 24 hours preceding the survey were categorized into seven food groups as (1) grains, roots, and tubers, (2) legumes and nuts, (3) milk and milk products, (4) flesh foods, (5) eggs, (6) vitamin-A rich fruits and vegetables, and (7) other fruits and vegetables. Finally, the food groups consumed by the child were added together to obtain dietary diversity score [17].

#### Food insecurity

Food insecurity was measured by HFIAS (Household Food Insecurity Access Scale) tool developed by FANTA (Food and Nutrition Technical Assistant) project. The tool has nine questions asking household’s about the three domains of food insecurity: feeling uncertainty of food supply, insufficient quality of food, and insufficient food intake and its physical consequences in the last month. The households participating in the study were categorized into the four levels of food-security (food secure, mildly food insecure, moderately food insecure and severely food insecure) based on the guideline’s recommendation [18].

#### Wealth index

Household wealth index was constructed using household asset data through PCA (Principal Component Analysis) based on interview responses adopted from Ethiopian Demographic and Health Survey. The presence or absence of each household items such as plow oxen, table, Animal-drawn cart, chair, etc. were asked and their responses were coded as ‘0’ for No and ‘1’ for Yes. Finally, the common factor score for each household was produced for grouping households as lower, middle and higher wealth quantile households [19].

#### Under-nutrition

chronic energy deficiency (malnutrition) was assessed using WHO guideline. The WHO Anthro 2005 software was used to calculate Z-score. MUACZ cut-off-point of negative two (−2) was used to define under-nutrition[20].

#### Data management and analysis

Data were entered into Epi-Data software version 1.4.4.0 and analyzed with Stata software version 14 (College Station, Texas).Proportions, means and standard deviations (SD) were used to describe the study population by independent variables and anemia. Bivariate logistic regression was done using thirty independent variables to identify the candidate variables (p-value < 0.25) for multivariable regression. Finally, predictors of anemia were determined using multivariable logistic regression model among the selected eleven candidate variables. Multi co-linearity, interaction and mediation among independent variables were checked based on the assumptions such as tolerance, variance inflation factors (VIF), correlation coefficient of interaction, and others to assure the fitness of logistic regression model‥ The independent variables used for checking multi co-linearity, interaction and mediation were; age of child, age of mothers,, religion, occupation, family size, meal frequency of mother, meal frequency of child, initiation of complementary feeding, receiving anti-helmenthic drug, sex of child, dietary diversity score of child (DDS) and household food insecurity access scale (HFIAS). The statistical significance was determined at p-value<0.05 at 95% confidence interval.

#### Quality control

Interviewers and laboratory technicians were trained for two days prior to data collection. A pilot study was done among 50 children who were selected from areas outside the actual study area. The data collection was regularly supervised by trained supervisors and the investigators.

#### Ethical consideration

The actual data collection was started after Wolaita Sodo University College of health sciences and medicine approved the study. Local administrative bodies (Damot Gale, Boloso Bombe, Humbo and Sodo Zuria district health offices) were also communicated and provided permission to undertake the study. Finally, written informed consent was obtained from the mothers/caregivers. Children diagnosed with anemia were counseled and those with hemoglobin concentration below 9.0 g/dl were referred to health facility for further treatment and follow up.

## Result

### Socio-demography

A total of 990 children were included in the survey making 99.7% response rate. The mean age of children was 14.96 months with a standard deviation of 5.37 months. Around 434(43.8%) of the children were females; 293(29.6%) were born with short birth interval (less than 2 years interval); 191(19.3%) were born from young mothers (mothers aged 15-24 years);and 60 (6.1%) were raised by single parents (Table 1).

**Table 1:**
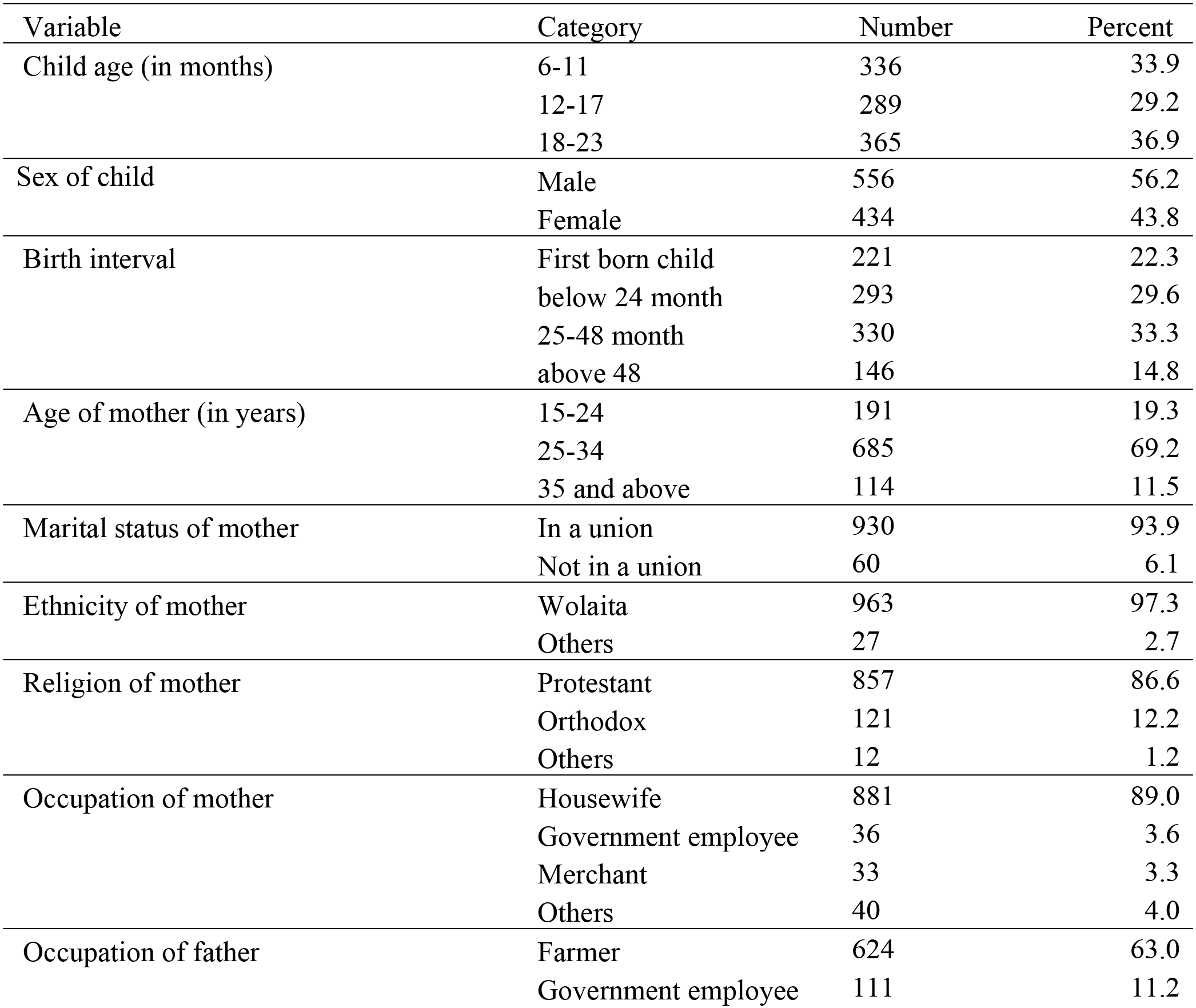
Socio-demographic characteristics of children aged 6-23 months residing in rural districts of Wolaita zone, 2016 (N=990)

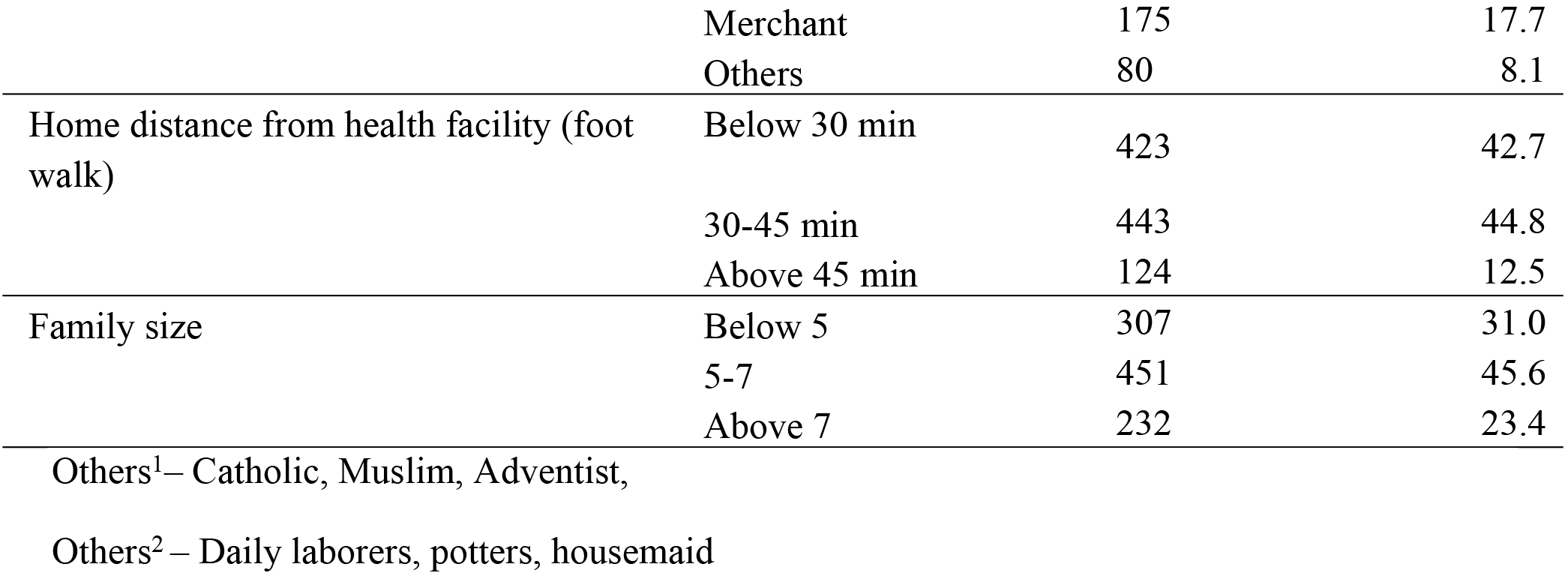

### Health and nutrition characteristics of children

The majority (93.1%) of children was fully immunized and only 11% of them were given anti-helminthic drugs to treat intestinal parasites. A total of 409 (41.3%) children had at least one of the following symptoms within the past two weeks preceding the survey: cough, difficulty of breathing, fever, diarrhea (with/without blood), or visible parasite on stool. Based on the Midupper arm circumference (MUAC) measurement, 688(69.5%)of the children were well-nourished, whereas the rest 302(30.5%) had chronic energy deficiency.

The children had mean dietary diversity score of 3.4 with 1.5 standard deviation, and only 422 (42.6%) of them consumed more than half of the recommended seven food groups. Furthermore, more than one-third (36.3%) drank either tea or coffee in their usual meal (that contained tannins which reduce iron absorption). Nearly one out of ten children stopped breastfeeding at least one week before the survey (Table 2).

**Table 2:**
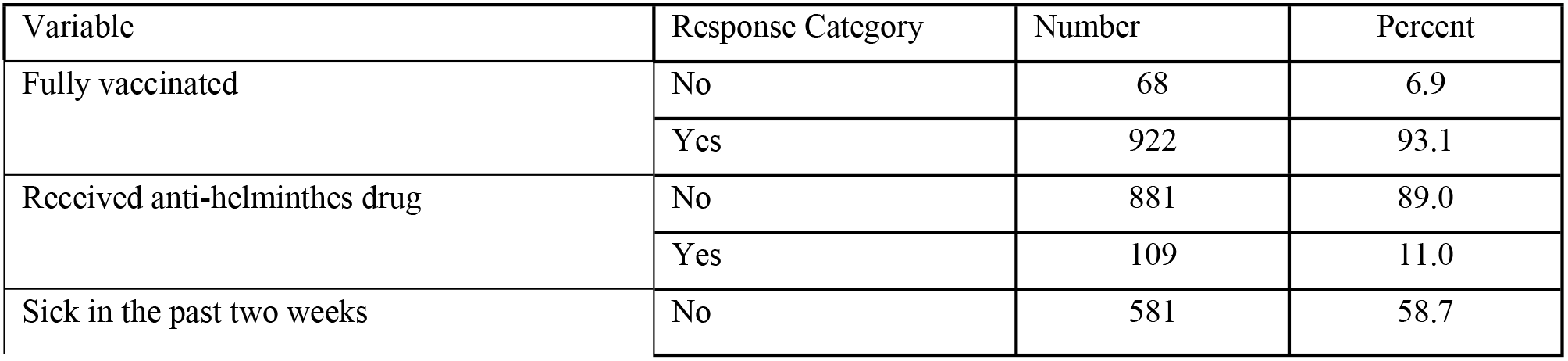
Health and nutrition characteristics of children in rural districts of Wolaita Zone, 2016 (N=990)

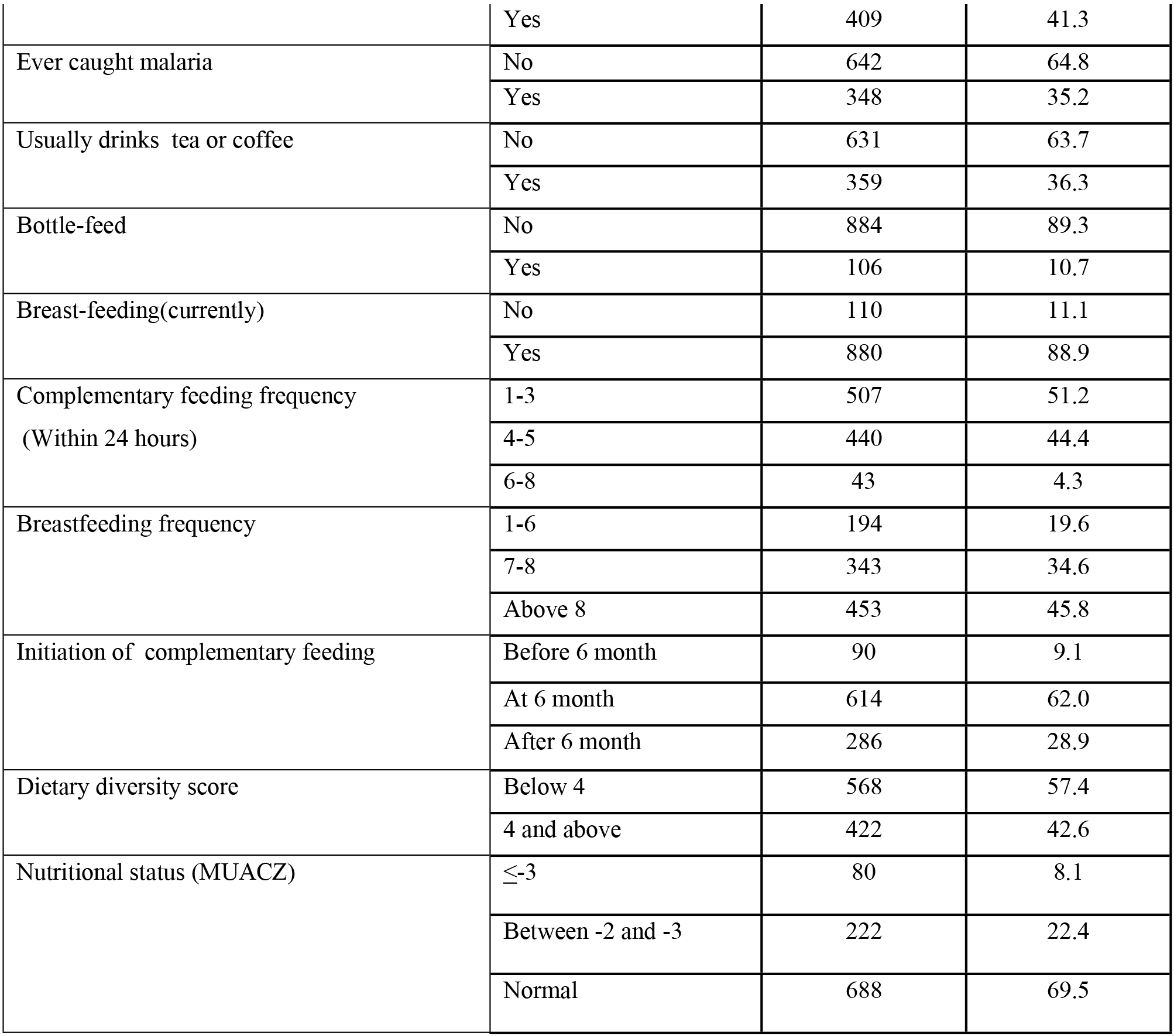

Based on the household food insecurity access scale measurement (HFIAS), 628 (63.4%) of the children were born and lived in food-secure households; whereas the rest were born and lived in a household suffering from food insecurity. Four hundred and four (40.8%) mothers fully attended the four WHO recommended ANC visits, while only 86(8.7%) of them had no ANC visit. The prevalence of maternal anemia was 13.4%; and three-fourth (734) of them usually consumed fewer than four meals per day (Table 3).

**Table 3:** Maternal health and nutrition in rural districts of Wolaita Zone, 2016 (N=990)

| Variable | Category | Number | Percent |
| --- | --- | --- | --- |
| Household food insecurity | Food secure | 628 | 63.4 |
|  | Mildly food insecure | 78 | 7.9 |
|  | Moderately food insecure | 208 | 21.0 |
|  | Severely food insecure | 76 | 7.7 |
| Number of pregnancies | 1-2 | 397 | 40.1 |
|  | 3-4 | 298 | 30.1 |
|  | Above 4 | 295 | 29.8 |
| Antenatal care follow-up | No visit | 86 | 8.7 |
|  | 1-3 visit | 500 | 50.5 |
|  | 4 and above | 404 | 40.8 |
| Place of delivery | Home | 446 | 45.1 |
|  | Health facility | 544 | 54.9 |
| Maternal anemia | Yes | 133 | 13.4 |
|  | No | 857 | 86.6 |
| Frequency of meals/day | Less than or equal to 3 | 734 | 74.1 |
|  | Above 3 | 256 | 25.9 |

### Magnitude and severity of anemia

The mean hemoglobin concentration of the children was 10.44 g/dl (95% C.I: 10.36, 10.52 g/dl) with standard deviation of 1.30 g/dl, and with a minimum and maximum hemoglobin concentration of 6.64 to 17.31g/dl. Overall, 650 (65.7%) of the children had anemia with hemoglobin level below 11g/dl (95% C.I: 62.6% - 68.6%). Most of them were mildly (10-10.9g/dl) and moderately (7-9.9g/dl) anemic accounting for 278 (28.1%) and 368 (37.2%), respectively; whereas only 4 (0.4%) of children were severely anemic with hemoglobin level below 7g/dl (Figure 1). The prevalence of anemia decreased as age increased i.e. from as high as 72.6% among infants aged 6-11 months to as low as 61.4% among young children aged 18-23 months (Figure 1).

**Figure 1:**
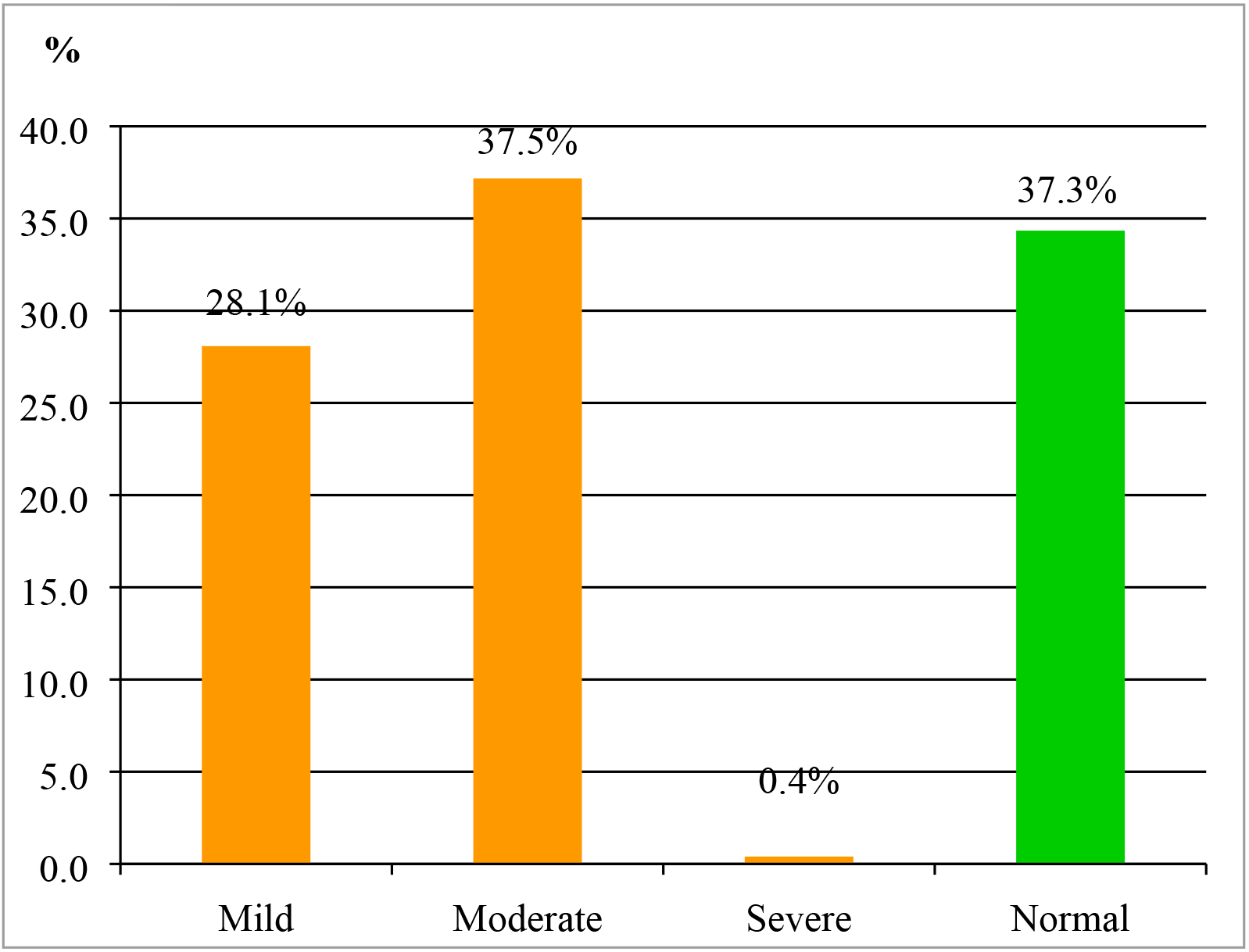
Magnitude and severity of anemia among children aged 6-23 months in rural districts of Wolaita Zone, 2016 (N=990)

### Factors associated with anemia

Bivariate logistic regression showed that age of child, age of mother, religion, occupation of mother, family size, child’s meal frequency, initiation of complementary feeding, receiving antihelminthic drugs, sex of child and food insecurity were associated with anemia in children (P<0.05).

However, in the multivariable analysis age of child, religion, family size and initiation of complementary feeding time were not associated with child anemia (p>0.05); whereas maternal age and occupation, child’s anti-helminthic drug intake, sex, dietary diversity score, and household food insecurity were identified as predictors of child anemia in the multivariable analysis (p<0.05).

The multivariable logistic regression showed that children of mothers whose age was 35 years and above were two times more likely to be anemic as compared to children of mothers whose age was between 15-24 years, AOR=1.96 (95% C.I: 1.01, 3.85), p=0.049. Children of government employee mothers had 61% lower chance of being anemic as compared to children of housewife mothers, AOR=0.29 (95% C.I: 0.09, 0.47), p=<0.001. Similarly, children of merchant mothers had 57% lower chance of getting anemia than children of housewife mothers, AOR= 0.43 (95%C.I: 0.20, 0.92), p= 0.029. In contrary, children of mothers whose occupation was classified as‘other occupation’(daily laborers, potters and housemaid) had greater chance of being anemic than children of housewives, AOR= 3.17 (95%C.I: 1.35, 7.43), p= 0.008.

Receiving anti-helminthic drugs was found to be inversely related with childhood anemia. Accordingly, children who received anti-helminthic drugs had 61% lower chance of getting anemia than their counterparts, AOR= 0.39 (95% C.I: 0.24, 0.63), p= <0.001. Female children are also twice more likely to be anemic than male children, AOR= 1.76 (95% C.I: 1.30, 2.38), p= < 0.001. Nutritional characteristics are also found to be predictive factors of childhood anemia. Accordingly, children who consumed poor dietary diversity (DDS < 4)were found to be more likely to be diagnosed anemic than their counterparts, AOR=1.40 (95% C.I: 1.03, 1.92), p= 0.034. Last but not least, children from moderately food insecure households were two times more likely to be anemic as compared to those from food secure households, AOR=1.72 (95% C.I: 1.07, 2.32), p= 0.023 (Table 4).

**Table 4:**
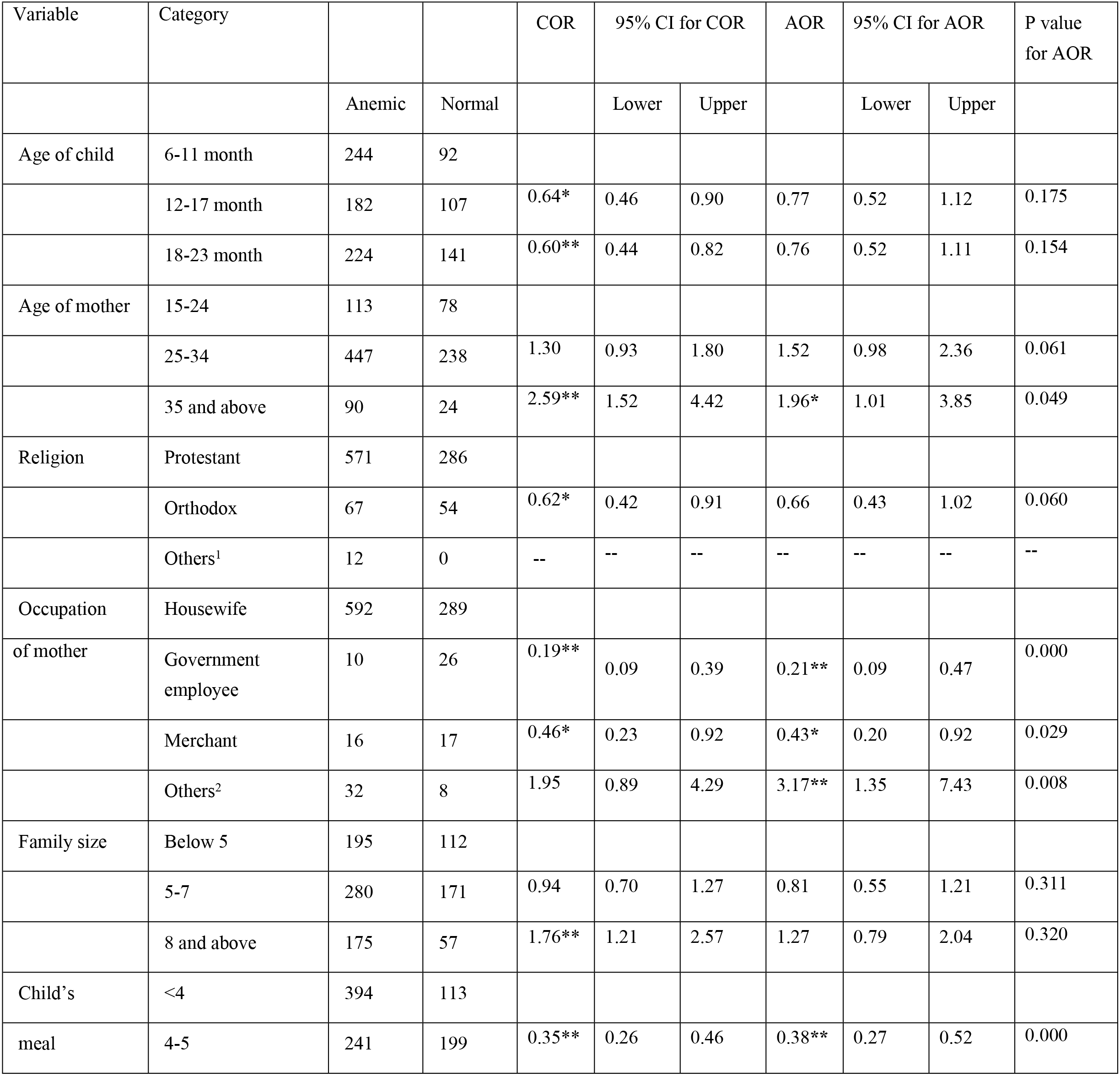
Multivariable analysis of factors associated with anemia among children in rural districts of Wolaita zone, 2016 (N=990)

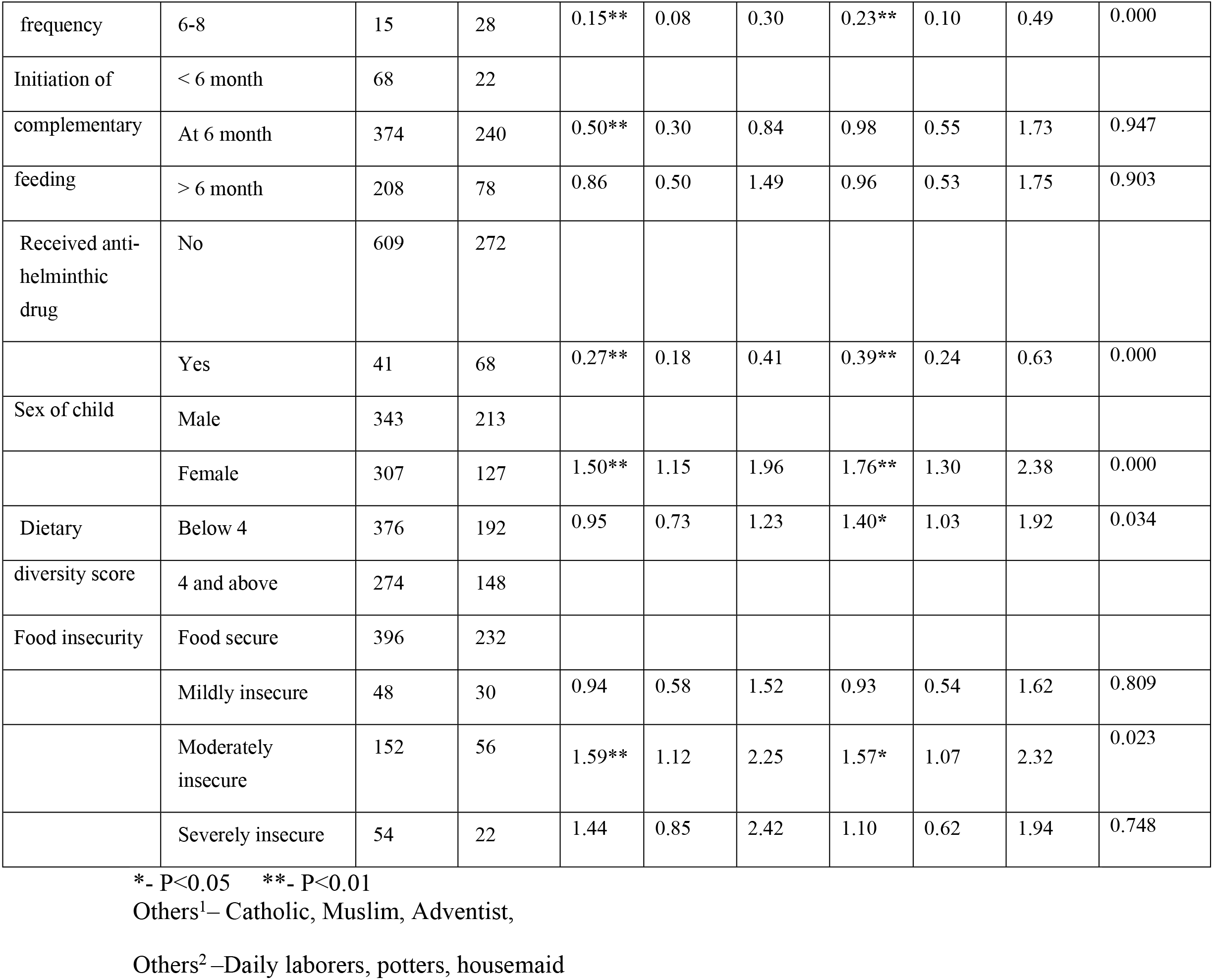

## Discussion

Anemia remains one of the major health problems that result in grave health outcomes in developing countries despite progresses seen in nutrition interventions. The Global Burden of Diseases study [21] shows that anemia in children is one of the most common causes of child death in Ethiopia, and continues to be a major public health problem. Similarly, our study showed that two-third of the young children in rural Wolaita had anemia standing as a severe public health problem (above the WHO cut-off point of 40%) [22]. On the other hand, studies elsewhere in Ethiopia [23] reported similar findings although severe anemia was found to be lower in the study area (0.4%) as compared to Ethiopia’s national rate among young children (3.5%) [12].

UNICEF and WHO recommend adequate breastfeeding, iron supplementation and fortification, and nutrition education for mothers [24] in order to curve health loss (high morbidity and mortality) due to iron deficiency anemia in children. Ethiopia’s nutrition strategy puts weight on the above nutrition intervention with the vision of ensuring adequate micronutrient intake for all children and its citizens [25]. Implementation of the national nutrition strategy with focus to young children will be vital in preventing and treating anemia in rural Wolaita.

The present study showed that maternal age, type of occupation, child’s anti-helminthic drug intake, sex of child, dietary diversity score and household food insecurity were associated with child anemia.

Studies have shown that consuming diverse food prevents anemia although this is a difficult option for households in developing countries where there is recurrent food insecurity problem. In conformity to this study, several studies [23, 26, 27] have shown children who had poor dietary diversity score had a higher chance of having anemia. Similarly, WHO report has shown that feeding children diverse foods increases the bioavailability of micronutrients including iron, and this is one of the recommended practices [28]. For instance, eating iron rich animal source food items such as flesh meat, organ meat, poultry; and non-animal source food items such as legumes and green leafy vegetables increase the bioavailability of iron and enables micronutrient demands for children. Consumption of fruits, vegetables, and tubers that are good sources of vitamins A and C, and folic acid enhances the absorption and utilization of iron [24].

De-worming intestinal parasites had also a positive impact on prevention of anemia among children [24]. Similarly, in this study, children in rural Wolaita who had undergone de-worming had lower odds of having iron deficiency anemia. A report of study conducted in 25 Sub-Saharan countries stated that de-worming and iron intake for more than six months prevented anemia in children[15]. De-worming is an essential public health intervention as intestinal parasites especially hookworm infection results in intestinal blood loss which in turn, contributes to anemia.

Household food security had also a significant role on preventing anemia among children [24]. In this study, we found that food insecurity was associated with the development of anemia, which was higher among children of moderately food insecure households. Studies elsewhere showed similar finding [29], and it is important to address food insecurity problems in rural areasto prevent anemia in children of such areas. In the absence of such measures, it will be difficult to achieve global and national targets of micronutrient problems. The efforts to ensure and assist local people with food security may need implementation of programs funded by government and non-governmental organizations. This could be an important means of intervention if such programs are implemented according to the standard [24] and prioritize people with severe or moderate food insecurity problems. Nevertheless, this study did not attempt to identify households supported by such programs.

Sex of the child was also one of the factors which influenced blood hemoglobin level of children. Many studies found the correlation between sex and hemoglobin level [29, 30, 31, 32]. According to those studies, the role of sex varies from place to place which gives different gender value for the sex of child with respect to the culture of the community. In the present study, female children were more likely to have anemia than males, although precise cause could not be spelled out. Some of the reasons could be high prevalence of sex bias (negatively affecting the feminine gender), late initiation of complementary feeding for female children and the community’s belief of greater need of care for male children than females [33]. Based on the finding of this study and other similar studies, female children are not favored to get iron rich food, which increases the risk of iron deficiency anemia. Therefore, it is important to educate the rural community about the importance of providing iron rich food for all children regardless of sex. Further studies may be required to validate the role of gender in iron deficiency anemia in Ethiopian context.

This study also showed that maternal age and occupation were correlated with children’s hemoglobin level. The risk of children’s anemia increased with the age of their mothers. In line with this finding, a study in Northeastern Brazil also showed similar trends of iron deficiency anemia [34]. On the other hand, parental occupation was also one of significant predictors of children’s hemoglobin level [27]. In this study, mothers who made relatively better incomes i.e. government employee and merchant mothers had a lower chance of having anemic children as compared to housewives. This indicates that anemia is significantly prevalent among families with less educated parents and low income, indicating the role of social inequality in the development of anemia [35]. Therefore, it will be important to ensure women with lower income either providing them opportunities to make more income or supplemental food for their children.

## Conclusion and recommendation

Based on the World Health Organization cut-off point, anemia is found to be a severe public health problem among young children residing in rural districts of Wolaita zone. Nearly two third of children aged 6-23 months were diagnosed to be anemic having hemoglobin level below 11g/dl. Poor dietary diversity, female sex of child, failure to take anti-helminthic drug, household food insecurity, maternal age and occupation were significantly associated with child anemia.

A large majority of children in the rural Wolaita were anemic, and the need for proven public health intervention such as food diversification, provision of anti-helminthic drugs and ensuring household food security is crucial. In addition, nutrition education and diet diversification through provision of alternative sources of income for women might be useful interventions.

## List of abbreviations

ANC: Antenatal Care
AOR: Adjusted Odds Ratio
C.I: Confidence Interval
COR: Crude Odds Ratio
EDHS: Ethiopian Demographic and Health Survey
FANTA: Food and nutrition technical assistant
Hb: Hemoglobin
HFIAS: Household Food Insecurity Access Scale
MUACZ: Mid Upper Arm Circumference Z-score
PCA: Principal Component Analysis
SD: Standard Deviation
SNNPR: Southern Nations, Nationalities and Peoples Region
UNICEF: United Nations Children’s Fund
WHO: World Health Organization

## Declarations

### Ethical approval and consent to participate

The actual data collection was started after Wolaita Sodo University College of Health Sciences and Medicine approved the study. Local administrative bodies were also communicated about the study and permission was obtained ahead of the study. Finally, informed written consent was obtained from the mothers/caregivers. Children diagnosed with anemia were counseled and those with hemoglobin concentration below 9.0 g/dl were referred to health facility for further treatment and follow up.

### Consent for publication

Consent to publish this manuscript from the participants was deemed not applicable since the manuscript does not contain any individual person data.

### Availability of data and material

The datasets used and analyzed during the current study are available from the corresponding author on reasonable request.

### Authors’ contribution

MA: conceived the study and designed the experiments. MA and MM: obtained ethical clearance, and supervised data collection. MA, BA and BY Performed statistical analysis, interpreted the data and revised the article. All authors read and approved the final manuscript.

## Acknowledgements

We would like to acknowledge Wolaita Sodo University for financial support for field work. We also extend our gratitude to respective district administrative bodies, data collectors, data clerk and study participants for their cooperation and technical assistance.

## Authors’ Information

MA has Masters of Public Health (MPH) in reproductive Health specialty and works at Wolaita Sodo University as assistant professor. MM has Masters of Public Health (MPH) in reproductive Health specialty and works at Wolaita Sodo University as assistant professor. BA has Masters of Science (MSc) in Biomedical Sciences and works as Associate professor at Wolaita Sodo University. BY has PhD in Public Health, and he is a Visiting Scientist at the Department of Global Health and Population, Harvard T.H Chan School of Public Health.

